# A comparative study of navigation behaviours in ants

**DOI:** 10.1101/2024.10.24.619962

**Authors:** Xianhui Shen, Antoine Wystrach, Uriel Gélin, Tristan Charles-Dominique, Kyle W. Tomlinson

**Affiliations:** Center for Integrative Conservation & Yunnan Key Laboratory for Conservation of Tropical Rainforests and Asian Elephants, Xishuangbanna Tropical Botanical Garden, Chinese Academy of Sciences, Menglun, Mengla 666303, Yunnan, China; University of Chinese Academy of Sciences, No. 19A Yuquan Road, Beijing 100049, China; Centre de Recherches sur la Cognition Animale, Centre de Biologie Intégrative, Université Paul Sabatier, CNRS, Toulouse, France; Centre for Biodiversity Dynamics in a Changing World (BIOCHANGE), Section of Ecoinformatics and Biodiversity, Department of Biology, Aarhus University, Aarhus, Denmark; AMAP, Univ Montpellier, CIRAD, CNRS, INRAE, IRD, Montpellier, France; Centre National de la Recherche Scientifique (CNRS), Sorbonne University, Paris, France

**Keywords:** canopy cover, foraging strategy, morphological traits, environmental cues, olfaction

## Abstract

Ants inhabit a vast range of ecosystems and exhibit wide morphology. They are expert navigators employing a handful of well-understood navigational strategies. However, the specific relationships among ant navigation behaviours, ecology, and morphology remain unclear, highlighting the need for comparative studies across diverse species. Here, we conducted field displacement experiments with 15 ant species across different habitats, assessing the prevalence of path integration, view-based navigation, olfactory trail following, and backtracking. We further tested whether use of particular navigation strategies was correlated with variation in morphological traits that could affect navigation efficiency, namely body size, eye size (view-based, path integration) and scape length (olfactory). There was a negative correlation between path integration and olfaction across different ant species, and no other clear trade-offs were identified between navigational strategies. Olfactory navigation emerged as the most dominant strategy. Path integration was also prevalent but limited to arboreal ants. View-based navigation was observed in both ground-foraging and tree-climbing ant species, and, unexpectedly, backtracking was also widespread. Species with larger eyes and body size showed a stronger preference for view-based navigation. However, no significant relationship was found between eye size or antennal scape length with preference for either path integration or olfaction. These results highlight the diversity and specialization of navigational strategies in ants, which appear to depend on the species’ ecological niche and morphological traits. Our study confirmed that path integration performs better in open sky environments, while view-based navigation appears more effective in cluttered habitats. We also showed the importance of plasticity in both foraging strategies and navigational profile at individual and colony levels, demonstrating the adaptability of ants’ navigation strategies to their environment.

## Introduction

How to navigate in the world is a task faced by most animals, making navigation an inherently comparative field. We have a good idea of the general evolution of navigation strategies across the big branches of the animal kingdom: from early taxis behaviours, such as kinesis in unicellular (Peng et al., 2016), taxis towards food sources in fly larvae (Chappell et al., 2022; Heaton et al., 2018), to long-range migrations in the monarch butterflies and birds [4], as well as the use of map-like representation in mammals(Cheng & Press, 2012; C. A. Freas & Cheng, 2022). But how specifically navigation behaviours are related to the ecology or morphology of species remains unclear and requires comparative approaches across phylogenetically more closely related species, sharing a similar navigational task and morphological design.

Ants are ideally suited for this endeavour. First, there is a vast number of species exhibiting substantial morphological diversity and inhabiting a wide variety of ecosystems (Drager et al., 2023; Hölldobler & Wilson, 1990; Parr et al., 2017). Second, ants are all excellent navigators that share a common navigational task: central place foraging, which involves searching for food and bringing it back to the nest (Huber & Knaden, 2015; Lach et al., 2010). Third, the navigational toolkit of ants is composed of a handful of well-understood strategies, the reliance on which varies across species and can be tested through simple experiments directly in the field (Graham & Mangan, 2015; Wehner, 2020). These include path integration, view-based landmark navigation, olfactory navigation, and other backup strategies such as systemic searching and backtracking (Wehner, 2020).

First, path integration, a vector navigation system, enables foragers to continuously keep an estimate of the straight-line distance and compass direction of their nest by continually integrating their own displacement. The information of compass direction is mainly based on celestial cues (the position of the sun, polarized light, as well as other light gradients) (Heinze et al., 2018; Müller & Wehner, 1988; Wystrach et al., 2014) and odometer records (stride integrator and self-induced optic flow) (Heinze et al., 2018; Ronacher & Wehner, 1995; Wittlinger et al., 2006).

Second, view-based navigation allows ants to navigate through the recognition of terrestrial cues, such as landmarks, surrounding panorama, skyline, and canopy patterns (C. Freas & Spetch, 2022; Graham & Cheng, 2009; Hölldobler, 1980; Macquart et al., 2006; Schwarz et al., 2014; Wystrach, Beugnon, et al., 2011). Information of direction is obtained by comparing the current view to visual memories obtained during the previous trips (Wystrach et al., 2012; Wystrach, Schwarz, et al., 2011; Zeil, 2012, 2023).

Third, olfactory-based navigation is a strategy employed by socially foraging ants. guiding recruits to food sources through pheromone trails, which can take various forms (Chalissery et al., 2022; Graham & Cheng, 2009; Jackson & Ratnieks, 2006; Wystrach, Beugnon, et al., 2011).

Fourth, the backup strategies, systematic searching and backtracking, increase the chance of foragers to find their goal when lost. These typically express when the other strategies provide no information (C. Freas et al., 2019; Wystrach et al., 2013). In systematic searching, the foragers perform loops of increasing size, starting from and returning close to a central area where the target location is likely to be found (Schultheiss & Cheng, 2011; Wehner & Srinivasan, 1981). During backtracking, ant foragers reverse their usual goal-heading direction, moving opposite to the homing route they recently travelled, which increases their chances of returning to familiar terrain (Wystrach et al., 2013).

The mechanisms underlying these navigational strategies have been studied in-depth in very few model species, and our current understanding of variation across species remains very limited, and mostly based on anecdotal evidence. Notably, studies on solitary foraging ant species dominate the field (Deeti et al., 2020; C. Freas & Spetch, 2019; Heinze et al., 2018; Müller & Wehner, 1988, 2010; Schwarz et al., 2020; Wehner, 1987, 2019; Wystrach, Beugnon, et al., 2011; Wystrach, Schwarz, et al., 2011). The apparent dominance of view-based navigation skills among those ant species (which is the reason why they have been chosen as models) suggest that they are favored as a consequence of their solitary foraging lifestyle, and notably their lack of clear olfactory cues such as pheromone trails. However, studies on ant species using pheromone trails (group or mass recruitment foraging) also showed their ability to use individual strategies such as path integration (C. Freas et al., 2019; C. A. Freas et al., 2019) or view-based navigation (Buehlmann & Graham, 2022; Fourcassie & Beugnon, 1988; Fukushi, 2001; Notomi et al., 2022). So, whether and how the foraging ecology of ants impacts their navigational strategies remains unclear.

Habitat structure may be a critical determinant of navigation strategies. Ants living in bare habitats, such as salt-pans, where terrestrial visual cues are minimal and celestial information is rich, such as *Cataglyphis fortis or Melophorus sp.,* favour path integration for navigation (Cheng et al., 2014; Schultheiss et al., 2016; Wehner, 2003). Ants inhabiting environments richer in visual landmarks but still with plenty of celestial cues, such as in semi-arid deserts (*Melophorus bagoti or Cataglyphis velox*) use both view-based navigation and path integration (Kohler & Wehner, 2005; E. L. G. Legge et al., 2014; Mangan & Webb, 2012; Wystrach et al., 2015). Finally, some forests dwellers (*Gigantiops destructor* and *Myrmecia croslandi*) are none to excel in view-based navigation and rely very little on path integration (Beugnon et al., 2005; C. A. Freas & Cheng, 2022; Macquart et al., 2006; Murray et al., 2019; Narendra, Gourmaud, et al., 2013). However, this evidence remains sparse and inconclusive, as it is merely based on a few solitary foraging species and the observed differences also co-vary with other ecological traits (e.g. thermophilic scavenger vs. visual predator) (Buehlmann et al., 2011).

Ants span huge differences in size and morphology, from a few millimeters (e.g. *Linepithema*) up to 3 cm (e.g. *Myrmecia*) and morphological traits are expected to impact navigational strategies. Regarding vision, the resolution and the visual field of compound eyes in ants are influenced by the number, area and arrangement of ommatidia (Land, 1997). Eye structure is loosely associated with ecological traits across species (Jelley & Barden, 2021) but can vary strongly with the individual’s cast within species (Narendra et al., 2010). Bigger ant workers have larger area eyes with greater numbers of ommatidia than conspecific smaller workers (Baker & Ma, 2006; Schwarz et al., 2010; Zollikofer et al., 1995), and ants with larger compound eyes have better visual acuity, but whether this influence their reliance on visual cues for navigation, such as landmarks and/or celestial cues is unclear (P N & Narendra, 2018). Regarding olfaction, antennal sensilla play a crucial role in chemoreception that is essential for olfactory navigation (Schneider, 2003; Zube et al., 2008). The macroglomeruli in the antennal lobe can exist and are typically involved in processing social information such as trail-pheromone (Kelber et al., 2009). Variation in antenna length may affect ability to detect chemical trails by increasing the surface area for chemical receptors, but this is associated to reliance over olfactory cues in the context of navigation is unclear. Overall, whether and how size and other morphological traits affect navigational behaviour remains largely unknown and dedicated comparative studies testing linkage between morphology and navigational behaviour are needed.

In this study, we assessed the reliance over of the four main navigational strategies (path integration, view-based navigation, olfactory navigation and backtracking) across a large number of ant species, sampled across a range of canopy covers. The species used exhibit substantial morphological differences, inhabit contrasting environments, and belong to different clades (15 species, 11 genera, and 5 subfamilies). We investigated the influence of canopy coverage on navigation strategy prevalence, as canopies coverage can be viewed as a proxy to estimate the prevalence of celestial and terrestrial cues. Those comparisons were conducted at both species level and within species level for a subset of widely distributed ant species, to test whether vegetation structure selected among navigation strategies, and whether ant species showed flexibility in employing navigational strategies across canopy cover gradients. We further tested whether navigational strategies were linked to specific morphological functional traits and foraging behaviour metrics.

Our predictions were as follows: (1) Solitary foraging ants are more likely to employ path integration or view-based navigation, while socially foraging ants may prefer olfactory trails. (2) As canopy cover increases, ants would shift their navigational preference from path integration to view-based navigation or olfaction, both within and across species (3) We anticipate a strategic trade-off between path integration and either view-based navigation or olfaction across species, as increased canopy cover would lead ants to prefer olfactory or view-based navigation. (4) Ants with larger eyes are predicted to favor path integration and view-based navigation, while those with longer antennal scapes are more reliant on olfactory trails.

## Materials and methods

### Study sites

Our experiments were conducted from May to November 2020 and from March to July 2021 in two sites. The first site was in Xishuangbanna Tropical Botanical Garden (XTBG), Chinese Academy of Sciences, located at Menglun, Mengla, Yunnan, China (101.4167°E, 21.6833°N), and experiences a tropical monsoon climate. This site covers an extensive area of 1,125 hectares, encompassing tropical rainforest and planted gardens. The rainforest has dense canopy coverage and complex understory plants, while plant gardens have simpler compositions and lower canopy coverage. A pre-investigation at XTBG revealed a large abundance and a rich diversity of ant species (around 150 species). The second site, located in, Hutiaoxia (HTX), Shangrila, Yunnan, China (100.1365°E, 27.2342°N), experiences a cold temperate monsoon climate. The area hosts three vegetation types: shrubland, broad-leaf forest fragments, and pine woodlands. A pre-investigation conducted in this region identified approximately 30 species. This variation provided an opportunity to explore the relationships between navigational strategies at inter- and intra-species levels across different vegetation types. Within XTBG, experiments were performed in five artificial landscapes: the flower garden (ACC: 0.43), the bamboo garden (ACC: 0.79), the fruit garden (ACC: 0.78), scientific research center (ACC: 0.72), and rainforest edges (ACC: 0.77). In Hutiaoxia, experiments were carried out in the three vegetation types: shrubland (ACC: 0.36), pine woodland (ACC: 0.81), and broad-leaf forest (ACC: 0.66).

### Study species

A total of 48 colonies were tested, including 15 species, 11 genera, and 5 subfamilies (Table S1). Within this sample, 23 colonies were tested in XTBG, encompassing one colony each of *Dolichoderus thoracicus*, *Paratrechina sp.1*, *Pheidole elongicephala*, *Pheidole plagiaria* and *Tetramorium kheperra*, two colonies each of *Aphaenogaster sp.3*, *Camponotus nicobarensis*, *Polyrhachis dives* and *Tetraponera rufonigra*, three colonies of *Camponotus parius* and seven colonies of *Odontoponora denticulata*. In HTX, 25 colonies were tested, including one colony each of *Aphaenogaster sp.2* and *Myrmica sp.1*, eleven colonies of *Formica japonica* and twelve colonies of *Aphaenogaster sp.1*. The last two species were found in all three vegetation types at the site.

### Experimental procedure

#### (1) Searching for Candidate Ant Nests

We offered various food items (tuna, egg cake, and fresh deceased termites) to foragers on the ground in the various sub-sites. Subsequently, we tracked the foragers who had picked up a food item back to their nests and marked those nests as potential candidates for the study.

#### (2) Foraging Recruitment

We marked a circle with a radius of 30 cm using one rope centered around the entrance of the candidate nest. These circles functioned as goniometers segmented into 24 sections marked by numbers along the ropes (Figure 1). To establish a spatial reference, the 0° of the goniometers were aligned with the northern direction using a magnetic compass. To ensure unimpeded travel for foragers, the circle rope was elevated by using twigs and stones. For arboreal ants, which established nests on tree trunk or bamboo, the base of the plant served as the center of the goniometer.

**Figure 1.**
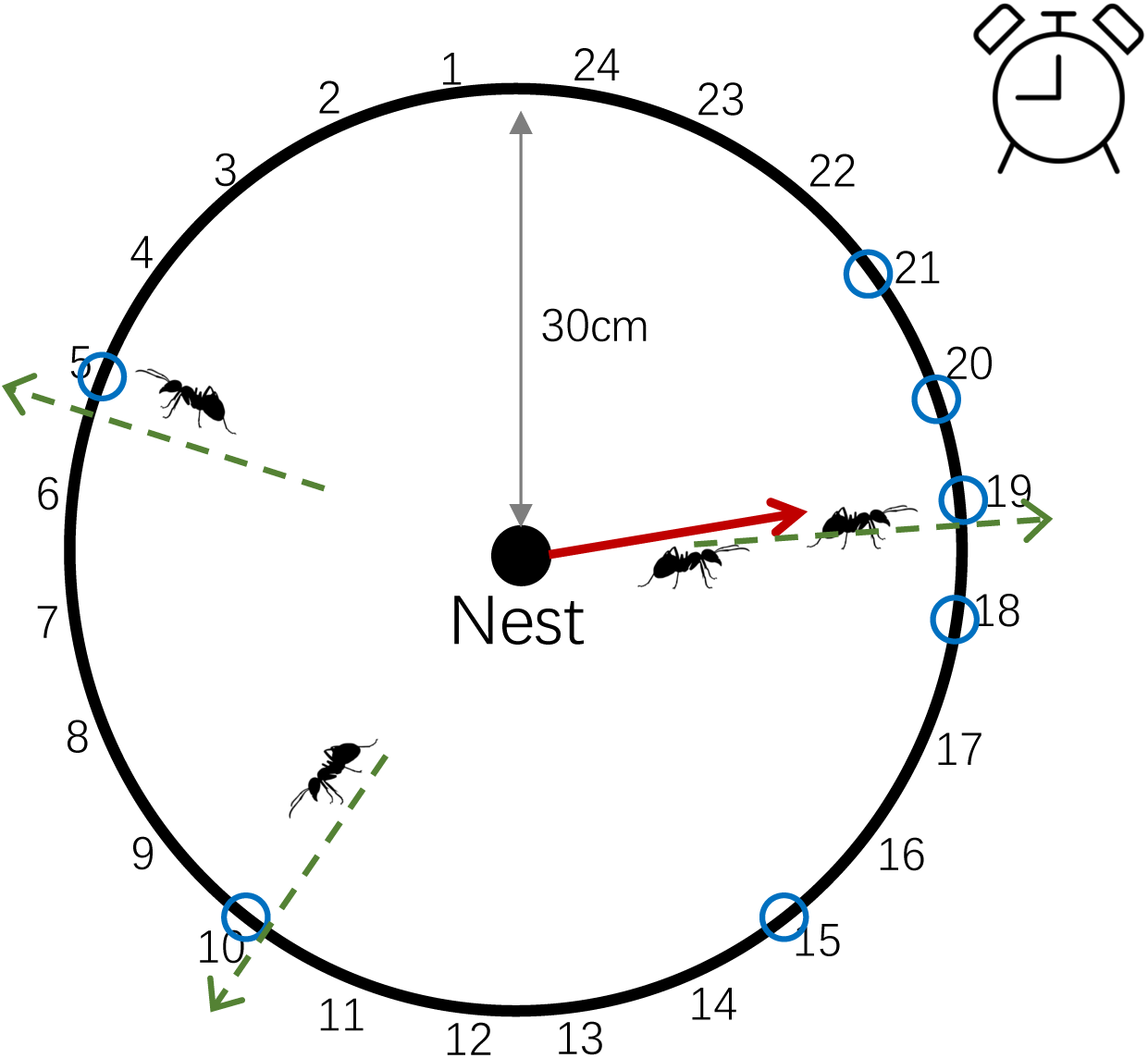
Measuring foraging activity and Foraging Directional Concentration (foraging R) based on the outbound heading of foragers. The heading directions of 20 outbound foragers (green dashed arrows and blue circles) were recorded as they crossed the 30 cm distance. The resultant vector length (foraging R, the red solid arrow) indicates how concentrated the foragers exit directions are. The time taken for 20 foragers to exit serve to calculate an index of foraging activity (foraging activity (ants/min) = 20/recorded duration).

We observed the outbound travel of foragers from the candidate nest and placed a feeding site along prevalent foraging direction, positioned at least two meters away from the nest entrance. Then, two foraging recruitment metrics (Traniello, 1989) (Table S2)were recorded: (1) Foraging Activity: the rate (ants/min) based on the first 20 foragers to come out from the nest and cross the 30 cm distance marked by the goniometer (Foraging Activity (ants/min) = 20/total_recorded_duration). This metric served as a proxy of foraging strategies, with colonies performing solitary behaviour characterized by smaller foraging activity values, while mass recruitment displayed larger values, and group recruitment fell in the middle. (2) Foraging Directional Concentration (foraging R): The heading directions of 20 outbound foragers were recorded 30 cm from the nest entrance using the goniometer, and we computed the mean resultant vector length (R) of the circular distribution (Batschelet, 1981). This value (foraging R) ranges from 0 (scattered) to 1 (concentrated) and indicates how concentrated the foragers’ outbound heading direction around the nest was. Solitary foraging individuals are expected to produce small values, because their foraging directions are independent, whereas ants following group and mass recruitment tend to produce high values, because most foragers go toward a same food source.

#### (3) Navigational strategy

We used displacement experiments to investigate the reliance on different navigational strategies. We used two identical wooden boards and one red transparent plastic sheet (Figure 2). Each wooden board and the red sheet featured a goniometer drawn on its surface, comprising a 60 cm-diameter circle segmented into 24 sections, each representing a 15° wedge. To maintain the wood boards horizontally on uneven ground, four stakes were installed. The red transparent plastic sheet was fixed on an iron frame with four stakes (10 cm high). During these displacement experiments, the 0° of the goniometers on both wooden boards and the red transparent plastic sheet were consistently aligned with magnetic north using a magnetic compass.

**Figure 2.**
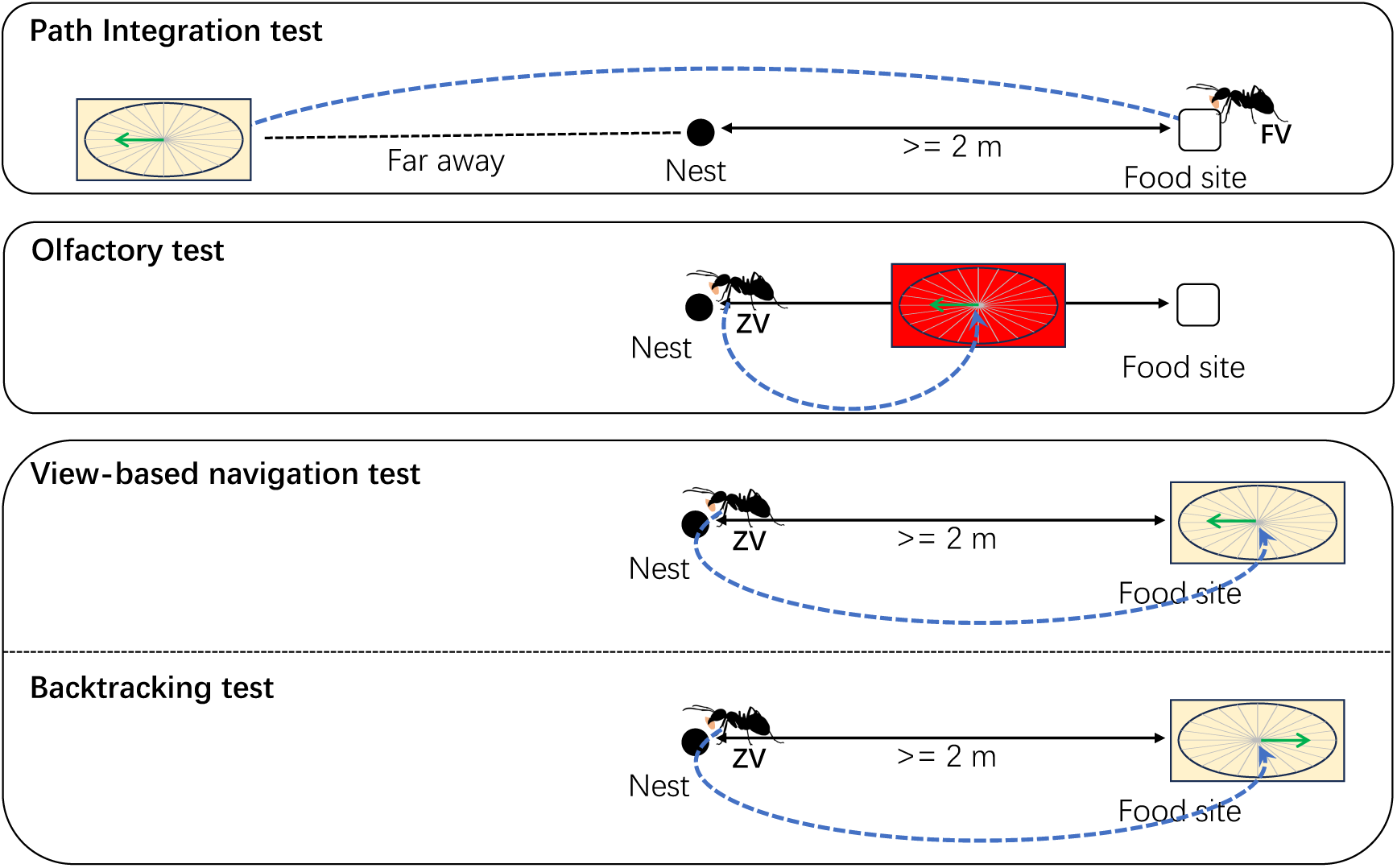
Scheme of displacement experimental setups achieved with homing ants. Homing ants trained to a feeder (open square) located at least 2m away from the nest (black dot) where captured with food in their mandibles in a tube, displaced (broken blue arrow lines) and released on a wooden goniometer (yellow square with sector) where their heading at distance of 30 cm was recorded. The yellow squares with goniometers represent the wooden boards for displacement, while the red square with a goniometer represents the red transparent plastic sheet used for the olfactory test. The green arrows indicate the theoretical direction of the strategy, the blue broken lines with arrows indicate the passive displacement effected in the dark tube between the capture and release points. “FV” (full vector ants) and “ZV” (zero vector ants) indicates the path integration state when released on the goniometer.

### Path Integration Test

Foragers at the feeder location that had just caught a food item and started their journey home were captured in a black tube. These are called full-vector ants (Knaden & Graham, 2016) as they possess a path integration homing vector pointing along the feeder-to-nest compass direction. These full vector ants were released in the center of the wooden board placed in a distant area away from the nest and thus unfamiliar to the ants, but where compass information was available for the ants to potentially express their path integration homing vector. The initial heading direction of individual foragers at the 30 cm goniometer was recorded, and we quantified whether the distribution was oriented in the feeder-to-nest compass direction.

### Olfactory Test

Homing zero-vector ants were captured just before entering their nest (as in the View-based Navigation test). These ants, so-called zero-vector ants no longer possess an informative path integration home vector. For the olfactory test, zero vector ants were released beneath the red transparent plastic sheet, positioned along the foraging route between the food source and the nest. The red transparent plastic sheet functioned as obstructing the visual information from landmarks or the sky, but still enabling the ant to follow olfactory marks on the floor. The individuals’ initial heading direction at 30 cm was recorded, and we quantified whether the distribution was oriented towards the nest direction

### View-based Navigation Test

Homing foragers that had just arrived at the nest entrance with a food item (zero-vector ants) (Wystrach et al., 2012) were captured in a black tube, transferred, and released on the wooden board placed 15 cm above the feeder location. This location provided a familiar view to the ants but prevented the use of olfactory cues. The individuals’ initial heading direction at 30 cm was recorded and we quantified whether the distribution was oriented towards the nest direction.

### Backtracking behaviour

was tested from the data obtained in the View-based Navigation test. Backtracking behaviour is observed in zero-vector ant species in the absence of familiar olfactory or visual cues, in which case the ants use compass cues to orient in the opposite, nest-to-feeder compass direction (C. Freas et al., 2019; Wystrach et al., 2013). Therefore, we quantified whether the distribution of ant headings in the View-based Navigation test was oriented in the direction opposite to the nest.

We aimed at testing 20 individual foragers for each of the path integration, view-based navigation, and olfactory experiments (Table S1).

### Morphological Traits

Three morphological functional traits that potentially correlate with the performance of navigational strategies were measured. Compound eyes are critical for detecting celestial cues in path integration and for view-based navigation (Schwarz et al., 2010; Wehner, 2020). Larger ant workers tend to have bigger eyes compared to their smaller conspecific workers (Baker & Ma, 2006). Therefore, body size of ants may influence the performance of navigational strategies. Antennae are essential for receiving olfactory information (Schneider, 2003), and the length of antennal scape may affect the ability to detect and process chemical information, thereby influencing reliance on using olfactory navigation. We measured these morphological functional traits based on protocols mentioned in Parr et al. (Parr et al., 2017) including: Weber’s length (ant size), used as a proxy of an ant body size; Relative eye size = π * (eye length/2) * (eye width/2)/ Weber’s length; Relative scape length = scape length/ Weber’s length.

For each species in every sub-site, morphological traits were measured. Six specimens were measured for monomorphic species (workers) and dimorphic species (minors), while ten specimens of workers were measured for polymorphic species, except for *Tetramorium kheperra*, which had only two workers measured. The limited sampling for *Tetramorium kheperra* was due to several days of heavy rain following the displacement experiment, which destroyed the nest and prevented further collection of specimens.

### Measuring Canopy Coverage

Using a CI-110 Plant Canopy Imager (CID Bio-Science), we measured the canopy coverage of each nest at ground level for three locations: the nest entrance (canopy_nest), the path integration test displacement point (canopy_pi), and the food resource site (canopy_food) (Table S1). The canopy coverage of these three points were highly correlated, canopy_nest with canopy_pi (r = 0.57, *p* = 0.029) and with canopy_food (r = 0.90, *p* < 0.001), and canopy_pi with canopy_food (r = 0.52, *p* = 0.049). Given these correlations, only canopy coverage of nest sites (canopy_nest) was used for further analysis, referred to as canopy cover hereafter.

### Data Analysis

Circular statistics (Batschelet, 1981) were conducted in MATLAB_R2022a. The analysis included the following steps:

#### Computation of a Score for each Navigational Strategy

For each tested nests, we computed a score for path integration (score_pi), view-based navigation (score_view), and olfaction (score_olf) (Table S2). For each test, we first calculated the mean resultant vector length and direction of the circular distribution of ants’ headings, and then measured the length of the orthogonal projection of this vector on the theoretical direction of the strategy (i.e., the feeder to nest compass direction). Thus, a score of 1 indicates that all tested ants were perfectly aligned along the strategy’s theoretical direction (Figure 2). A score of -1 indicates that all ants headed perfectly in the opposite direction, and a score of 0 indicates that ants were not biased along or against this theoretical direction (at the group level). All negative values were replaced by 0, resulting in a score varying from 0 to 1 and indicating how strongly oriented toward the strategies’ direction ants were.

We also computed a score for backtracking (score_bt) (Table S2). Backtracking was evaluated using the same test as the view-based navigation but in the opposite predicted direction (Figure 2). Consequently, the score_view and the score_bt can only be negatively correlated, a high score in view-based navigation implied a negative (and thus zero) score in backtracking, and vice versa. Because backtracking and view-based navigation are not independently estimated, we do not interpret correlations between backtracking and view-based navigation. Note that this approach yields us to underestimate the prevalence of Backtracking, as it can be observed only in species that did not rely on view-based navigation.

### Inter-species Analysis

#### (1) Correlation among navigational strategies across species

We calculated the correlation among the three navigational strategies scores across species in R. Mean scores of each navigational strategy for each species were first calculated to investigate species-level reliance on each strategy. The significant level for this study was set at 0.1.

#### (2) Factors explaining navigational strategies

Firstly, we checked the Spearman correlation among explanatory variables (foraging activity, foraging R, canopy cover), and morphological traits, to make sure the variables were not correlated with each other. We then performed RDA analysis to explore the relationship between navigational strategies (score_pi, score_view and score_olf) and explanatory variables. The significance of the RDA model was evaluated using a permutation test. Furthermore, a linear regression model was applied to the significant RDA model to investigate the correlation between the navigation strategy and the associated explanatory factors.

#### (3) Patterns of composition of navigational strategies in ants

We performed a cluster analysis across the 15 species using a heatmap based on the navigational strategies scores in R (ComplexHeatmap). These gave us four groups, for which we then compared against one another for scores of navigational strategies, morphological metrics, canopy coverage and foraging metrics. ANOVA and Kruskal-Wallis tests were used to test whether the grouping explained significant variation and Bonferroni correction were used to pairwise tests among the groups.

### Intra-species Analysis

We conducted intra-species analysis to investigate the flexibility of navigational strategies within species. This was achieved for the three species with multiple repeats from different colonies: *Aphaenogaster sp.1* (12 colonies), *Formica japonica* (11 colonies), and *Odontoponera denticulata* (7 colonies). To examine the reliance on specific navigational strategies, we used Levene’s test to assess the variance in scores between different strategies for each species. Additionally, we explored how variations in navigational strategy scores correlated with canopy cover, foraging activity and Foraging R variation across nests using Spearman correlation. Since the score_view for most colonies in *Aphaenogaster sp.1*, and *Odontoponera denticulata* was zero (Supplementary Table 1), it could artificially inflate correlation with other metrics, potentially leading to misleading results. Therefore, Levene’s test and Spearman correlations involving score_view was not calculated in *Aphaenogaster sp.1*, and *Odontoponera denticulate* to ensure unbiased results.

Variability in navigational strategies may be influenced by environmental factors (C. A. Freas & Schultheiss, 2018) and specific morphometric characteristics (Narendra, Alkaladi, et al., 2013; Schwarz et al., 2010). Specialization in a particular navigation strategy can possibly lead to trade-offs with their ability to perform other type of navigation strategies. To disentangle and quantify the importance of each factor, we used Spearman correlation to assess the strength of the relationship between these variables by combining and centering the data from three different species.

## Results

### Inter-species patterns

#### (1) Correlation among navigational strategies across species

Spearman correlation analysis revealed that there were no significant correlations among the navigation strategies, despite perhaps a tendency for a negative correlation between path integration and olfaction (r = -0.49, *p* = 0.070) (Table 1). The absence of general correlations suggests there are no universal trade-off or antagonistic constraints between the navigational strategies.

**Table 1.**
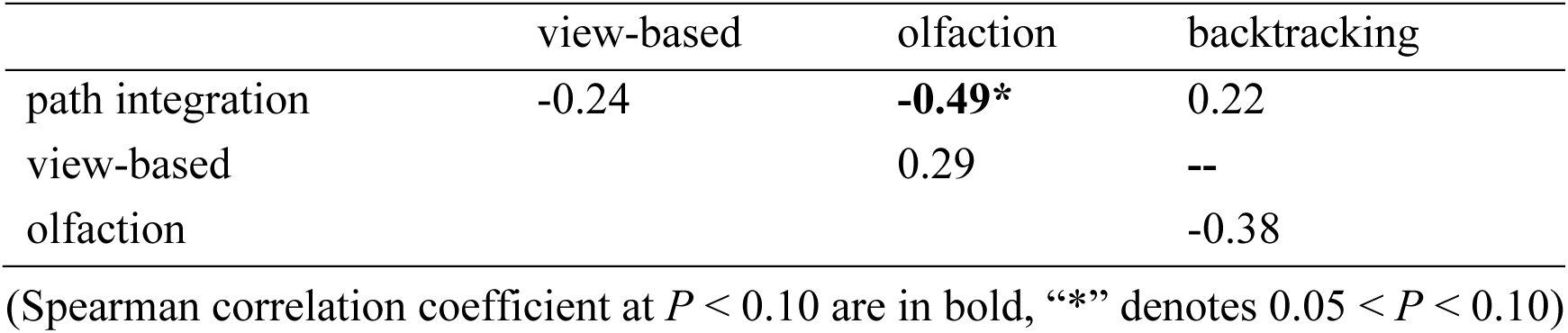
Spearman correlation among navigational strategies for all tested species.

#### (2) Predictors explaining navigational strategies

The results of Spearman correlation analysis revealed no significant correlation between foraging metrics (foraging activity and foraging R), morphometrics or canopy cover, except for positive correlation between ant size and relative eye size (Table 2).

**Table 2.** Spearman correlation coefficients among explanatory variables.

|  | foraging<br>R | Antsize | Relative<br>Eyesize | Relative<br>Scaplength | canopy<br>cover |
| --- | --- | --- | --- | --- | --- |
| foraging<br>activity | 0.11 | 0.15 | 0.37 | 0.21 | 0.09 |
| foraging R |  | -0.28 | -0.29 | 0.01 | -0.03 |
| Antsize |  |  | **0.62\*\*** | -0.31 | -0.31 |
| Relative<br>Eyesize |  |  |  | **-0.45\*** | -0.27 |
| Relative<br>Scaplength |  |  |  |  | 0.19 |

(Spearman correlation coefficient at $p < 0.10$ are in bold, “\*” denotes $0.05 < P < 0.10$ , “\*\*” denotes $0.01 < P < 0.05$ )

Results from the RDA analysis indicated that neither path integration nor olfaction displayed a relationship with foraging metrics (foraging activity and foraging R), morphological traits (ant size, relative eye size, relative scape length), or canopy coverage. On the other hand, the overall RDA model for view-based navigation (*p* = 0.069) and the first axis (RDA1) (*p* = 0.080) were suggestive. View-based navigation exhibited a significantly positive relationship with log-transformed relative eye size (ANOVA, F = 13.14, *p* = 0.0031, adj.R^2^ = 0.46) and ant size (ANOVA, F = 5.84, *p* = 0.031, adj.R^2^ = 0.26). As a consequence, absolute eye size displayed also a significantly positive relationship with view-based navigation (ANOVA, F = 12.62, *p* = 0.004, adj.R^2^ = 0.45) (Figure 3). Taken together, large ant size and large relative eye size implies large absolute eye size, suggesting the importance of big eyes for reliable use of view-based navigation.

**Figure 3.**
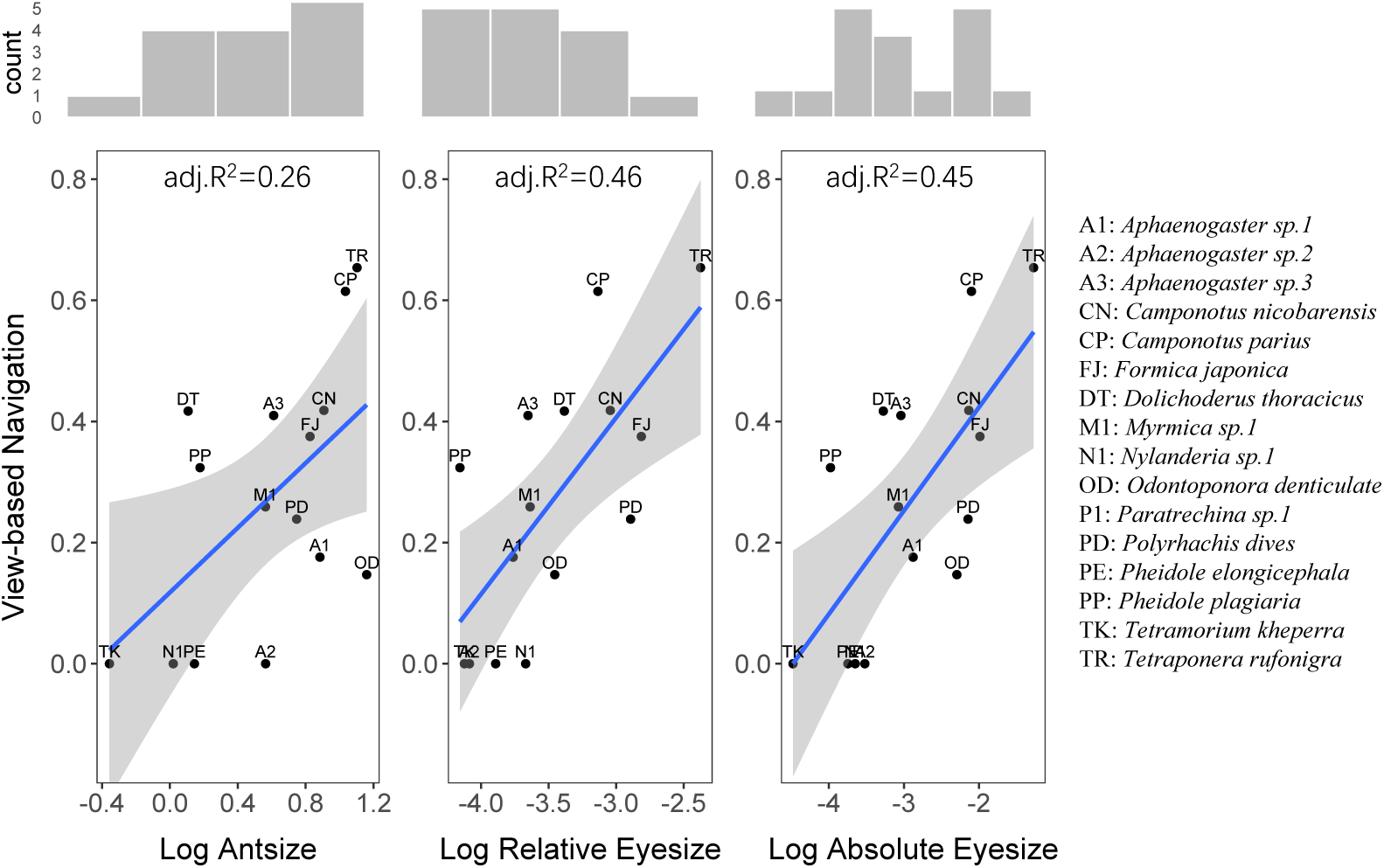
Relationship between morphological traits and view-based navigation (normalized transformed, score_view^(1/2)) among all tested ant species. The view-based navigation displayed significant positive relationship with (a) Log-transformed Antsize, (b) Log-transformed relative Eyesize, and (c) Log-transformed absolute Eyesize. The histograms on the top display the distribution of Log Antsize, Log Relative Eyesize and Log Absolute Eyesize.

#### (3) Composition of navigational strategies in ants

Fifteen ant species were clustered into four groups based on mean scores of path integration, view-based navigation, olfaction, and backtracking (Figure 4A). Group 1, including species of *Dolichoderus thoracicus*, *Camponotus nicobarensis* and *Polyrhachis dives,* exhibited the lowest score in path integration, high score in olfaction, low score in view-based navigation and no backtracking. Group 2, encompassing species of *Camponotus parius* and *Tetraponera rufonigra*, displayed medium score in path integration, high score in olfaction, the highest score in view-based navigation (and therefore backtracking ability cannot be determined). Group 3, including species of *Tetramorium kheperra*, *Odontoponora denticulata*, *Aphaenogaster sp.2*, *Paratrechina sp.1* and *Pheidole elongicephala*, exhibited high score in path integration, high score in olfaction, the lowest score in view-based navigation, but a high score in backtracking. Group 4, including *Formica japonica*, *Aphaenogaster sp.3, Pheidole plagiaria*, *Myrmica sp.1* and *Aphaenogaster sp.1*, demonstrated high score in path integration, high score in olfaction, low score in view-based navigation, and low score in backtracking.

**Figure 4.**
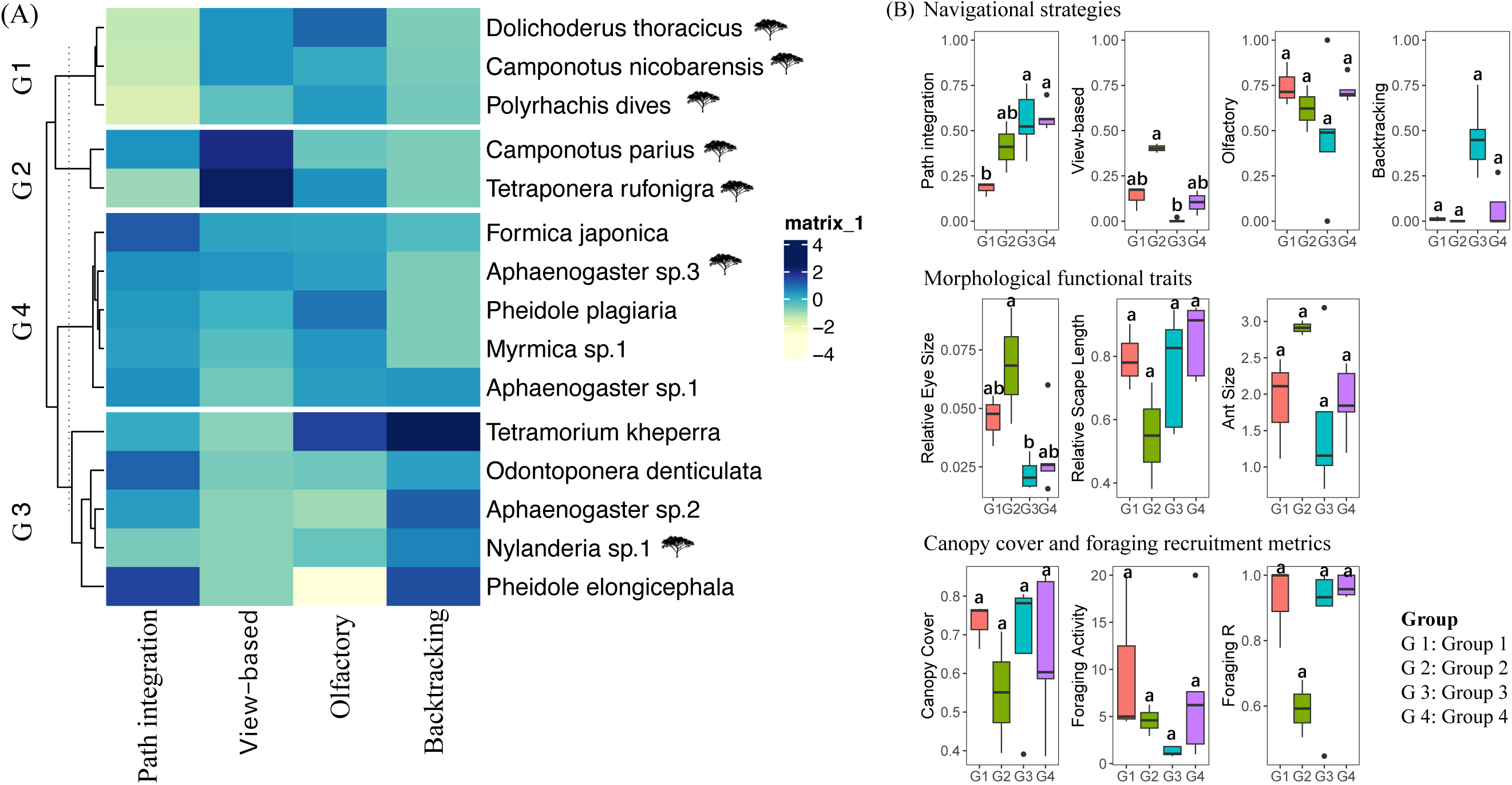
(A) Heatmap based on mean scores of navigational strategies, with 15 species were clustered into 4 groups. The species marked by a tree logo were observed to forage both on the ground and on trees or shrubs during the experiments. (B)The variation of scores of navigational strategies, morphological functional traits (relative eye size, relative scape length, ant size), canopy cover and foraging recruitment metrics (foraging activity and foraging R) of the four clustered groups. G1: Group 1, G2: Group 2, G3: Group 3, G4: Group 4.

The variation in scores of navigational strategies revealed that the score for olfaction was higher than that for path integration and view-based navigation across species (Figure 4B). Olfaction consistently emerged as the predominant and widely utilized (93%, 14 out of 15 species) navigational strategy, except in the case of *Pheidole elongicephala*, which notably lacked olfaction in cluster Group 3 (Figure 4A). Path integration was prevalent with high scores among species in Group 3 and 4, but was less frequently employed with lower scores in species of Group 1 and 2. Within Group 1, one colony of *Polyrhachis dives* demonstrated no utilization of path integration, as did a colony of *Tetraponera rufonigra* in Group 2 (Supplementary Table 1). Contrastingly, the scores for view-based navigation were almost consistently the lowest among the three navigational strategies, indicating a comparatively lesser reliance and low accuracy of view-based navigation across all but the two species forming Group 2. Species in Group 1 and Group 4 exhibited both low scores for view-based navigation and backtracking, meaning their orientation when released at the familiar food area (without path information) was rather random. Species in Group 3 displayed the lowest score for view-based navigation but the highest score for backtracking, indicating they headed in the direction opposite to their nest when released at the familiar feeder location without path integration information.

### Navigational plasticity within species

*Aphaenogaster sp.1* (12 colonies) and *Formica japonica* (11 colonies) were found in habitats with a wider range of canopy coverage (0.28 to 0.88) compared to *Odontoponera denticulata* (0.61 to 0.95), which were consistently found near the edge of relatively dense forests (Figure S1).

Navigational strategy scores varied greatly between colonies of the same species indicating plasticity in the utilization of navigational strategies (Figure 5). These three species exhibited rather low (all below 0.5), but variable scores for the view-based navigation, and a substantial proportion of colonies in each species – 9 out of 12 for *Aphaenogaster sp.1*, 5 out of 11 for *Formica japonica*, and 6 out of 7 for *Odontoponera denticulata* – scored 0. Conversely, these three species demonstrated relatively higher scores for olfaction and path integration, indicating a systematic preference for these strategies over view-based navigation. Path integration and olfaction scores did not differ between *Aphaenogaster sp.1* (F = 0.28, *p* = 0.60) and *Formica japonica* (F = 0.02, *p* = 0.89). However, in *Odontoponera denticulata*, the variance in scores for olfaction was higher than for path integration (F = 4.24, *p* = 0.062, Figure 5). This suggested a greater reliance on path integration over olfaction in *Odontoponera denticulata*.

**Figure 5.**
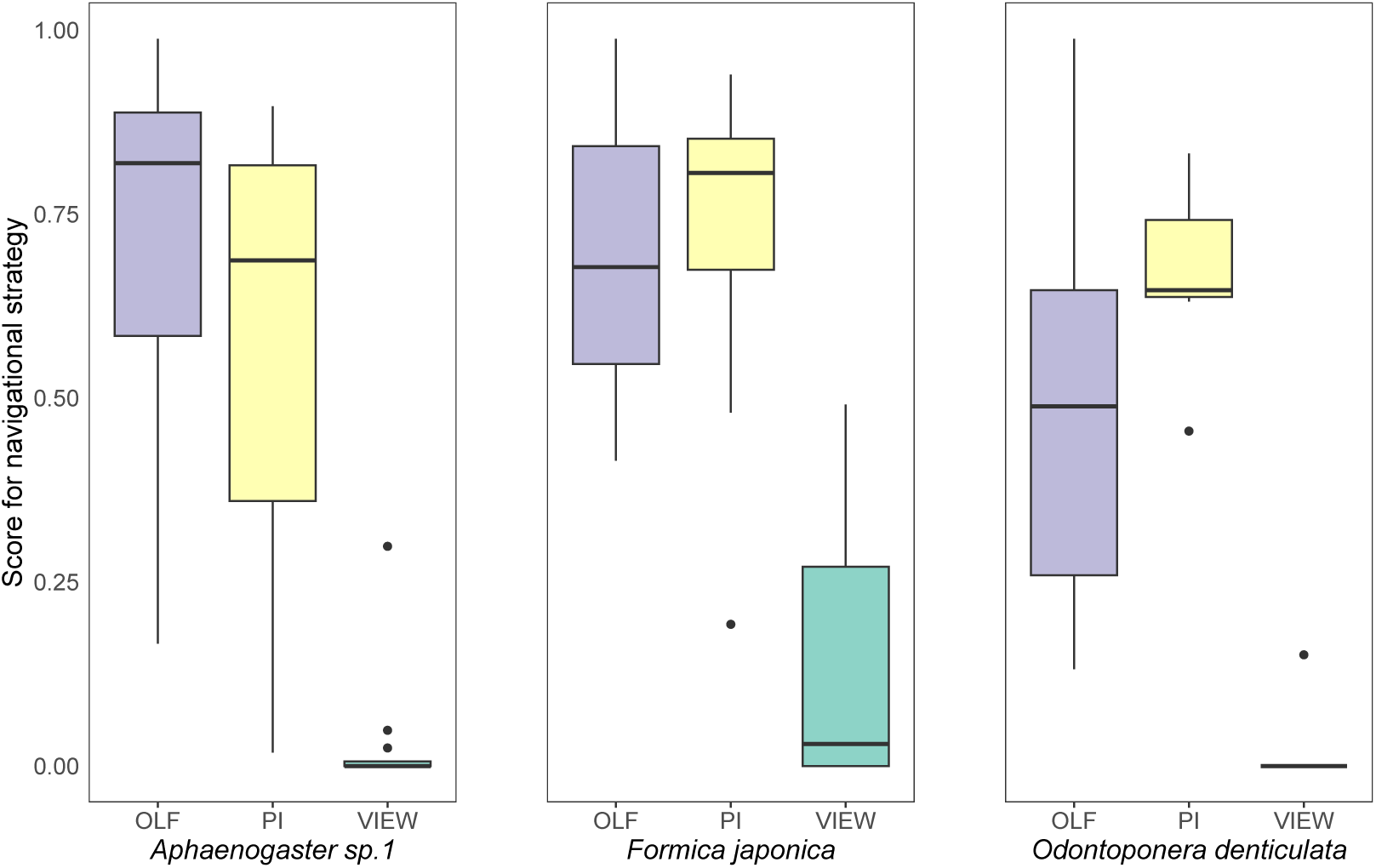
Scores of navigational strategies of *Aphaenogaster sp.1* (12 colonies), *Formica japonica* (11 colonies) and *Odontoponera denticulata* (7 colonies). OLF: olfaction, PI: path integration, and VIEW: view-based navigation.

Within species, no significant correlation was found between the navigational strategies (path integration, view-based navigation, or olfaction) and environmental or foraging metrics. However, several tendencies (p<0.1) appeared: in *Formica japonica*, canopy coverage exhibited a negative correlation with path integration (r = -0.54, *p* = 0.094), suggesting a preference for path integration in more open surroundings. For *Aphaenogaster sp.1*, canopy coverage showed a positive correlation with foraging R (r = 0.56, *p* = 0.060) and olfaction (r = 0.51, *p* = 0.087), olfaction exhibited a positive correlation with foraging R (r = 0.56, *p* = 0.063). Additionally, path integration positively correlated with foraging activity (r = 0.51, *p* = 0.093). These results suggest that in more closed canopies, *Aphaenogaster sp.1* increases recruitment and reliance on olfaction, and path integration. In *Odontoponera denticulata*, foraging activity positively correlated with path integration (r = 0.75, *p* = 0.066), suggesting that increased recruitment in *Odontoponera denticulata* corresponds to an increased reliance on path integration.

Analyzing these three species collectively, results revealed that the canopy coverage significantly correlated positively with view-based navigation (r = 0.43, *p* = 0.017), and tended to correlate negatively with path integration (r= -0.32, *p* = 0.083) (Table 4), suggesting that in more open canopy environments, ants rely more on path integration, while in more closed area, ants rely more on view-based navigation.

## Discussion

Our study presents a first comparative approach aimed at measuring and understanding the variation in the use of navigational strategies across ant species. Performance in the four navigational strategies tested here (Figure 4A) varied greatly across the 15 species studied. While the ant species were clustered into 4 groups based on their performance profile across our tests, the categorization was not distinctly clear. Variability in navigational strategy performance was not only observed between the groups but also within them (Figure 4A), and, across colonies of same species (Figure 5). Together, this highlights flexibility of navigational strategies employed by ant species both at the ultimate and proximal levels. Related to this, and perhaps surprisingly, no general correlation between the performance across our tests was observed, suggesting no universal trade-offs across the use of the different navigation strategies.

### Overestimated view-based navigation and underestimated olfaction

Contrary to its dominant role in the ant navigation literature (Buehlmann et al., 2020; C. A. Freas & Cheng, 2022; C. Freas & Spetch, 2022; Heinze et al., 2018), in our study, most species were bad at using view-based navigation (Figure 4B), that is, at deriving their goal direction from learnt familiar views even though our experimental situation facilitated the use of learnt views by releasing ants at their very familiar feeder location (Figure 2). Also, for the species that performed well, view-based navigation was never used as a sole strategy. The ability to use view-based navigation correlated strongly with ant size and eyes size (Figure 5). Indeed, bigger eyes enable higher resolution, and bigger ants can afford bigger eyes. This suggest that the general bias in the literature of investigating ants with large body size and eyes (for instance, ants from genus *Cataglyphis* species (Mangan & Webb, 2012; Schwarz et al., 2020), *Myrmecia* species (Islam et al., 2020; Narendra & Esquivel, 2017), *Melophorus bagoti* (Schwarz et al., 2014; Wystrach et al., 2012) and *Gigantiops destructor* (Beugnon et al., 2005; Macquart et al., 2006) may have led to a tacit overestimation of importance of view-based navigation by ants.

Inversely, the reliance of ants on olfaction has been probably underestimated in non-trail following ants (Buehlmann et al., 2020; Knaden & Graham, 2016). In our study, olfaction-based navigation prevailed for ant species across all different investigated ecological niches, even in apparently solitary foraging ant species such as *Odontoponera denticulata*. These findings suggest that if the use of view remains widespread like in their flying hymenopteran ancestor, ants, as walking animals, rely greatly on olfaction for navigation, except perhaps for a few ant species, which are typically the big with large eyes.

### Backtracking is widespread

Backtracking strategy, that is, the tendency to head opposite to the usual home compass direction when in the absence of other directional cues, which was shown in only three studies (C. Freas et al., 2019; Plowes et al., 2019; Wystrach et al., 2013), turned out to be surprisingly widespread. As a backup strategy, backtracking increases the chance of a lost individual to fall back on familiar cues (pheromone trials, nest odour, or familiar views). It relies on compass cues, likely celestial signals (Wystrach et al., 2013), but the mechanisms triggering this strategy seem to differ across the ant species. In the solitary foraging species *Melophorus bagoti*, backtracking is triggered when zero-vector foragers (i.e., devoid of directional information from Path integration) that have recently been exposed to the nest panorama, are released on unfamiliar terrain (Wystrach et al., 2013). For the socially foraging species, *Veromessor pergandei*, backtracking is triggered when homing foragers that have run along approximately 75%-85% of the pheromone column, are released off the column (C. Freas et al., 2019; Plowes et al., 2019). In another socially foraging species, *Formica obsuripes*, backtracking is elicited when foragers have completed over 50% of their homeward journey and encounter the absence of both familiar panorama view and pheromone cues (C. Freas & Spetch, 2023). In our study, the foragers consistently ran off their path integrator and/or the whole homing pheromone column until they reached their nest, but were then displaced at the feeder location, that is, within a visually *familiar* environment (2 meters from nests) (Figure 2). Therefore, backtracking could be observed here only in species that did *not* use view-based navigation (see Figure 4 (A) group 3), and it is thus possible that the species using view-based navigation would have equally shown backtracking behaviours if tested in visually unfamiliar environment. Thus, backtracking may have been underestimated in our study. The fact that we now have observed this strategy in three subfamilies – Myrmicinae (*Aphaenogaster sp.1* and *Aphaenogaster sp.2* of *Aphaenogaster* genus, *Tetramorium kheperra* of *Tetramorium* genus, and *Pheidole elongicephala* of *Pheidole* genus), Formicinae (*Nylanderia sp.1* of *Nylanderia* genus and *Formica japonica* of *Formica* genus) and Ponerinae (*Odontoponera denticulata* of *Odontoponera* genus) - suggests that backtracking may well be ancestral to ants. This calls for testing the existence of this strategy in other navigating insects.

### Navigational strategies and foraging strategies

The literature conveys the tacit assumption that the reliance of navigational strategies is largely dependent on the species foraging strategy, that is, whether the ants use pheromone trail for recruitment (and thus are thought to rely largely on olfaction) (Hölldobler et al., 2001; Jackson & Ratnieks, 2006) or forage solitary (and thus are thought to rely mostly on views or path integration) (E. Legge et al., 2014; Wystrach et al., 2012). The lack of clear correlations between olfaction and either path integration or view-based navigation in our analysis challenges this assumption. Furthermore, we observed great variation in foraging activity and recruitment metrics across colonies of the same species, and these correlated with reliance over different navigational strategies, highlighting the individual and colony-level plasticity in both foraging strategies and navigational profiles. For instance, olfaction correlated positively with foraging R (i.e., the concentration of departure directions) the across colonies of in *Aphaenogaster* sp.1. suggesting that the reliance over olfaction increases with the concentration of foragers for a single route, which may simply be explained by a higher concentration of pheromone trail. Also, path integration correlated positively with foraging activity across both *Aphaenogaster* sp.1 and *Odontoponera denticulata* (Table 3), which may be explained by a higher proportion of new naïve recruit, with little experience of the surrounding terrestrial cues and thus a higher reliance over celestial cues(Schwarz et al., 2017), to the food source. Overall, this highlights how navigational strategies can be tuned, at the proximal level, through the complex interplay between collective context and individual flexibility.

**Table 3.**
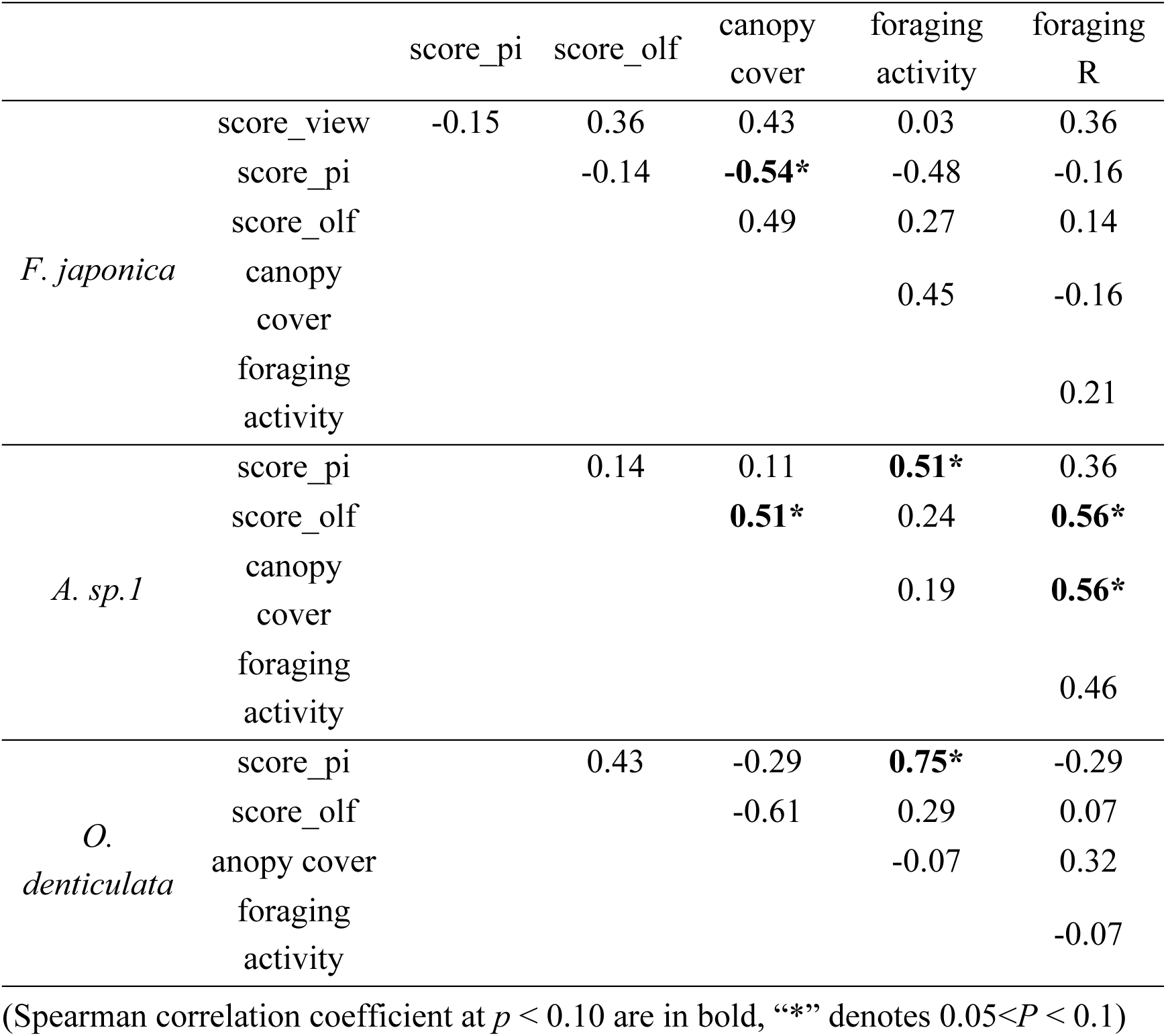
Spearman correlation coefficients among navigational strategies, canopy cover, foraging activity and foraging R of *Formica japonica*, *Aphaenogaster sp.1*, and *Odontoponera denticulata*.

### Influence of arboreality and canopy coverage on Path integration

Our results also confirm the general idea that path integration performs better in open sky environments –where celestial cues are plentiful and landmark are scarce – while view-based navigation excels in cluttered environments –where terrestrial visual information is plentiful and path integration errors may accumulate much quicker (Buehlmann et al., 2018; Wehner, 2020). This might be true not only across species, but also as a plastic response within species. Canopy cover was negatively correlated with path integration but positively correlated with view-based navigation across the three species, *Aphaenogaster sp.1*, *Formica japonica* and *Odontoponora denticulate* (Table 4). Within the species *Formica japonica*, canopy cover was negatively correlated with path integration, while within the species *Aphaenogaster sp.1*, canopy cover was positively correlated with view-based navigation (Table 3). However, despite this influence of the availability of celestial/terrestrial cues, our result showed no overall negative correlation between the reliance of path integration and view-based navigation, indicating no fundamental trade-off between the use of both strategies.

**Table 4.**
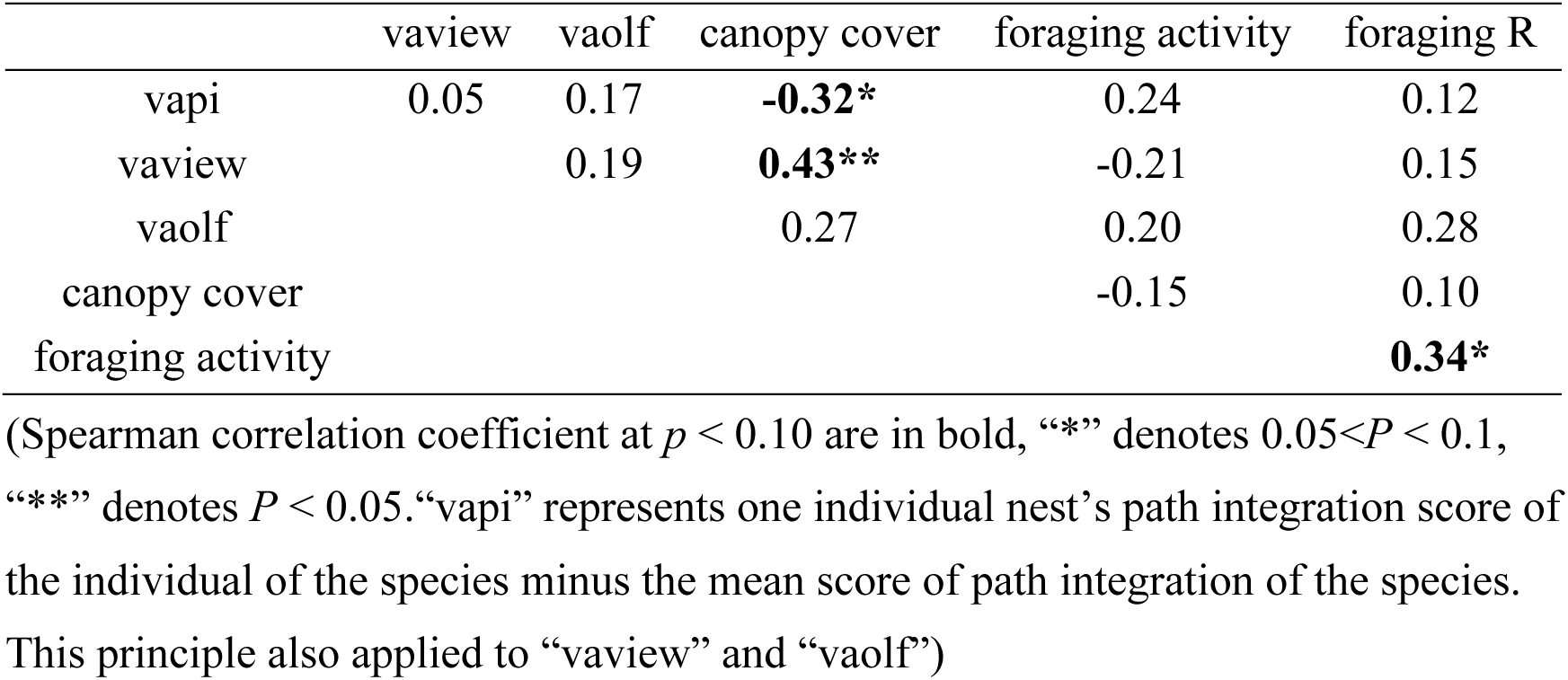
Spearman correlation coefficients among deviation from average of scores of three navigational strategies (path integration, view-based navigation, and olfaction), canopy cover and foraging activity.

|  | vaview | vaolf | canopy cover | foraging activity | foraging R |
| --- | --- | --- | --- | --- | --- |
| vapi | 0.05 | 0.17 | **-0.32\*** | 0.24 | 0.12 |
| vaview |  | 0.19 | **0.43\*\*** | -0.21 | 0.15 |
| vaolf |  |  | 0.27 | 0.20 | 0.28 |
| canopy cover |  |  |  | -0.15 | 0.10 |
| foraging activity |  |  |  |  | **0.34\*** |

(Spearman correlation coefficient at $p < 0.10$ are in bold, “\*” denotes $0.05 < P < 0.1$ , “\*\*” denotes $P < 0.05$ . “vapi” represents one individual nest’s path integration score of the individual of the species minus the mean score of path integration of the species. This principle also applied to “vaview” and “vaolf”)

Not all species relied on path integration, even though this mechanisms is definitely ancestral to the evolution of ants (Wehner & Srinivasan, 2003). Interestingly, the 5 out of 15 species that did not rely on path integration, were all observed to forage in trees (Figure 4A), whereas among the 10 other species that oriented well using path integration, only 2 were observed to forage in trees. Not relying on path integration when foraging in tree makes sense, not because odometric or compass errors may accumulate more quickly in trees, but simply because path integration is useful to guide new shortcut routes on the ground, but it is not effective in arboreal habitats as one cannot take a shortcut between two different branches. This idea corroborates previous research that has demonstrated that visual cues and pheromone cues are typically used for orientation in three-dimensional environments (Beugnon & Fourcassié, 1988; Fourcassie & Beugnon, 1988; C. A. Freas et al., 2018).

This comprehensive study revealed the flexibility of navigation strategies employed by ants by incorporating as many species as possible, highlighted the importance of plasticity in both foraging strategies and navigational profile at individual and colony levels, demonstrating the adaptability of navigation strategies employed by ants to their environment.

## Author contributions

A.W., U.G., T.C.D., and X.S. conceived and designed the project. X.S. carried out the fieldwork and data collection. X.S., A.W., U.G., T.C.D., and K.W.T. analysed data, wrote and revised the manuscript. K.W.T. got the funding.

## Acknowledgements

We thank Yun Lu, Dong Yang, Bo Li, Guangyou Zhang, Xiaqin Luo and Jianju Feng for field assistance.

