## Supplemental information for "A comparative study of navigation behaviours in ants"

**Supporting information**

**Article title: A comparative study of navigation behaviours in ants**

The following Supporting Information is available for this article:

**Table S1** Species, location, canopy cover and sample size for each tested ant nest.

**Table S2** Foraging recruitment metrics and scores of navigational strategies for each tested ant nest.

**Figure S1.** Canopy coverage of nest sites of *Aphaenogaster sp.1* (left), *Formica japonica* (middle) and *Odontoponera denticulata* (right).

**Table S1.** Species, location, canopy cover and sample size for each tested ant nest.

| Sample NO. | Species | Location | | Canopy cover | | | Sample size | | |
| --- | --- | --- | --- | --- | --- | --- | --- | --- | --- |
|  |  | site | sub-site | canopy_nest | canopy_food | canopy_pi | sample size_pi | sample size_view | sample size_olf |
| XTBG_NE_11 | *Camponotus parius* | XTBG | flower garden | 0.1900 | 0.3450 | 0.1830 | 20 | 20 | 20 |
| XTBG_NE_1 | *Camponotus parius* | XTBG | scientific research center | 0.5070 | 0.3380 | 0.4490 | 20 | 20 | 18 |
| XTBG_NE_32 | *Camponotus parius* | XTBG | scientific research center | 0.4850 | 0.3470 | 0.3600 | 20 | 20 | 20 |
| XTBG_NE_12 | *Camponotus nicobarensis* | XTBG | flower garden | 0.4930 | 0.5560 | 0.2680 | 20 | 20 | 20 |
| XTBG_NE_7 | *Camponotus nicobarensis* | XTBG | Bamboo garden | 0.8340 | 0.8230 | 0.8230 | 20 | 20 | 22 |
| XTBG_NE_3 | *Tetraponera rufonigra* | XTBG | fruit garden | 0.7980 | 0.7760 | 0.4790 | 20 | 20 | 20 |
| XTBG_NE_5 | *Tetraponera rufonigra* | XTBG | flower garden | 0.6180 | 0.7510 | 0.3250 | 20 | 20 | 20 |
| XTBG_NE_9 | *Polyrhachis dives* | XTBG | bamboo garden | 0.8020 | 0.7610 | 0.7740 | 20 | 20 | 20 |
| XTBG_NE_8 | *Polyrhachis dives* | XTBG | bamboo garden | 0.7240 | 0.8030 | 0.8170 | 20 | 20 | 20 |
| XTBG_NE_4 | *Dolichoderus thoracicus* | XTBG | fruit garden | 0.7670 | 0.8720 | 0.4790 | 19 | 27 | 16 |
| XTBG_NE_10 | *Odontoponera denticulata* | XTBG | scientific research center | 0.9480 | 0.8100 | 0.5270 | 16 | 16 | 16 |
| XTBG_NE_16 | *Odontoponera denticulata* | XTBG | rain forest edges | 0.7250 | 0.7560 | 0.5760 | 20 | 20 | 20 |
| XTBG_NE_19 | *Odontoponera denticulata* | XTBG | rain forest entrance | 0.6080 | 0.6230 | 0.5590 | 22 | 20 | 14 |
| XTBG_NE_20 | *Odontoponera denticulata* | XTBG | rain forest | 0.6090 | 0.5590 | 0.6480 | 20 | 20 | 20 |
| XTBG_NE_26 | *Odontoponera denticulata* | XTBG | rain forest entrance | 0.7670 | 0.7490 | 0.7640 | 22 | 20 | 20 |
| XTBG_NE_28 | *Odontoponera denticulata* | XTBG | rain forest | 0.9270 | 0.7490 | 0.9050 | 20 | 20 | 0 |
| XTBG_NE_29 | *Odontoponera denticulata* | XTBG | rain forest | 0.8880 | 0.7490 | 0.9440 | 21 | 20 | 7 |
| XTBG_NE_17 | *Aphaenogaster sp.3* | XTBG | rain forest entrance | 0.7360 | 0.7740 | 0.7380 | 20 | 20 | 20 |
| XTBG_NE_27 | *Aphaenogaster sp.3* | XTBG | rain forest | 0.9390 | 0.7490 | 0.9060 | 20 | 20 | 20 |
| XTBG_NE_31 | *Pheidole elongicephala* | XTBG | scientific research center | 0.6520 | 0.7140 | 0.7760 | 20 | 20 | 12 |
| XTBG_NE_38 | *Pheidole plagiaria* | XTBG | scientific research center | 0.8540 | 0.8100 | 0.8750 | 20 | 20 | 20 |
| XTBG_NE_36 | *Tetramorium kheperra* | XTBG | scientific research center | 0.8050 | 0.8190 | 0.6770 | 16 | 16 | 7 |
| XTBG_NE_37 | *Nylanderia sp.1* | XTBG | scientific research center | 0.7950 | 0.8550 | 0.7730 | 20 | 20 | 20 |
| HTX_NE_1 | *Formica japonica* | HTX | thicket | 0.4100 | 0.4330 | 0.3140 | 20 | 20 | 20 |
| HTX_NE_5 | *Formica japonica* | HTX | thicket | 0.3060 | 0.3400 | 0.2700 | 20 | 20 | 20 |
| HTX_NE_14 | *Formica japonica* | HTX | thicket | 0.3110 | 0.3190 | 0.3910 | 20 | 20 | 20 |
| HTX_NE_17 | *Formica japonica* | HTX | thicket | 0.3770 | 0.3250 | 0.7340 | 20 | 20 | 20 |
| HTX_NE_8 | *Formica japonica* | HTX | pine | 0.8430 | 0.8470 | 0.7760 | 14 | 15 | 15 |
| HTX_NE_30 | *Formica japonica* | HTX | pine | 0.7710 | 0.7670 | 0.8120 | 20 | 20 | 20 |
| HTX_NE_32 | *Formica japonica* | HTX | pine | 0.8820 | 0.8440 | 0.8060 | 20 | 20 | 20 |
| HTX_NE_18 | *Formica japonica* | HTX | broad-leaf fragment | 0.6910 | 0.7180 | 0.6120 | 20 | 20 | 20 |
| HTX_NE_21 | *Formica japonica* | HTX | broad-leaf fragment | 0.7410 | 0.7160 | 0.7390 | 20 | 20 | 20 |
| HTX_NE_22 | *Formica japonica* | HTX | broad-leaf fragment | 0.5290 | 0.6800 | 0.7100 | 20 | 20 | 20 |
| HTX_NE_27 | *Formica japonica* | HTX | broad-leaf fragment | 0.5900 | 0.6100 | 0.7610 | 20 | 20 | 20 |
| HTX_NE_2 | *Aphaenogaster sp.1* | HTX | thicket | 0.5420 | 0.3740 | 0.2890 | 20 | 20 | 20 |
| HTX_NE_4 | *Aphaenogaster sp.1* | HTX | thicket | 0.2820 | 0.2970 | 0.3270 | 20 | 20 | 20 |
| HTX_NE_15 | *Aphaenogaster sp.1* | HTX | thicket | 0.3260 | 0.3190 | 0.3250 | 20 | 20 | 20 |
| HTX_NE_16 | *Aphaenogaster sp.1* | HTX | thicket | 0.2990 | 0.3030 | 0.2620 | 10 | 10 | 10 |
| HTX_NE_6 | *Aphaenogaster sp.1* | HTX | pine | 0.8050 | 0.7950 | 0.8340 | 20 | 20 | 20 |
| HTX_NE_29 | *Aphaenogaster sp.1* | HTX | pine | 0.7820 | 0.7800 | 0.8100 | 20 | 20 | 20 |
| HTX_NE_31 | *Aphaenogaster sp.1* | HTX | pine | 0.8050 | 0.7950 | 0.8070 | 20 | 20 | 20 |
| HTX_NE_19 | *Aphaenogaster sp.1* | HTX | broad-leaf fragment | 0.7400 | 0.7130 | 0.6990 | 20 | 20 | 20 |
| HTX_NE_20 | *Aphaenogaster sp.1* | HTX | broad-leaf fragment | 0.6910 | 0.6780 | 0.6860 | 20 | 20 | 20 |
| HTX_NE_25 | *Aphaenogaster sp.1* | HTX | broad-leaf fragment | 0.5720 | 0.5970 | 0.7390 | 20 | 20 | 20 |
| HTX_NE_26 | *Aphaenogaster sp.1* | HTX | broad-leaf fragment | 0.7260 | 0.7410 | 0.7510 | 20 | 20 | 20 |
| HTX_NE_13 | *Aphaenogaster sp.1* | HTX | road side | 0.6660 | 0.6100 | 0.5550 | 20 | 20 | 20 |
| HTX_NE_28 | *Myrmica sp.1* | HTX | road side | 0.3860 | 0.3760 | 0.7720 | 20 | 20 | 20 |
| HTX_NE_3 | *Aphaenogaster sp.2* | HTX | thicket | 0.3910 | 0.3150 | 0.2710 | 21 | 18 | 14 |

**Table S2.** Foraging recruitment metrics and scores of navigational strategies for each tested ant nest.

| Sample NO. | Forage Activity | Foraging R | score_pi | score_olf | score_view | score_bt |
| --- | --- | --- | --- | --- | --- | --- |
| XTBG_NE_11 | 4 | 0.9684 | 0.4717 | 0.4551 | 0.0434 | 0 |
| XTBG_NE_1 | 15 | 0.5319 | 0.2556 | 0.8487 | 0.5275 | 0 |
| XTBG_NE_32 | 8 | 0.539 | 0.9282 | 0.1751 | 0.5633 | 0 |
| XTBG_NE_12 | 5 | 1 | 0.2018 | 0.3701 | 0 | 0.0159 |
| XTBG_NE_7 | 4 | 1 | 0.2019 | 0.9221 | 0.3499 | 0 |
| XTBG_NE_3 | 8 | 0.5967 | 0.5372 | 0.7007 | 0.4783 | 0 |
| XTBG_NE_5 | 2 | 0.4132 | 0 | 0.8015 | 0.377 | 0 |
| XTBG_NE_9 | 3 | 0.5586 | 0 | 0.7735 | 0.114 | 0 |
| XTBG_NE_8 | 6 | 0.9984 | 0.272 | 0.6547 | 0 | 0.055 |
| XTBG_NE_4 | 1 | 1 | 0.208 | 0.8791 | 0.174 | 0 |
| XTBG_NE_10 | 23 | 0.9606 | 0.7089 | 0.1317 | 0.1514 | 0 |
| XTBG_NE_16 | 24 | 0.9698 | 0.6325 | 0.3611 | 0 | 0.3052 |
| XTBG_NE_19 | 20 | 0.8036 | 0.7781 | 0.9903 | 0 | 0.3401 |
| XTBG_NE_20 | 16 | 0.9612 | 0.8341 | 0.596 | 0 | 0.3136 |
| XTBG_NE_26 | 38 | 0.8886 | 0.456 | 0.1586 | 0 | 0.2846 |
| XTBG_NE_28 | 12 | 0.9669 | 0.648 | 0.4896 | 0 | 0.0537 |
| XTBG_NE_29 | 31 | 0.9803 | 0.6448 | 0.6997 | 0 | 0.3831 |
| XTBG_NE_17 | 12 | 0.978 | 0.772 | 0.4748 | 0.0616 | 0 |
| XTBG_NE_27 | 8 | 0.8891 | 0.363 | 0.9089 | 0.2744 | 0 |
| XTBG_NE_31 | 11 | 0.9061 | 0.7604 | 0 | 0 | 0.5067 |
| XTBG_NE_38 | 1 | 1 | 0.5347 | 0.8371 | 0.1048 | 0 |
| XTBG_NE_36 | 25 | 0.9966 | 0.4808 | 1 | 0 | 0.7535 |
| XTBG_NE_37 | 11 | 0.4467 | 0.3319 | 0.5078 | 0 | 0.3411 |
| HTX_NE_1 | 10 | 0.9966 | 0.8423 | 0.5154 | 0.476 | 0 |
| HTX_NE_5 | 8 | 0.9902 | 0.8386 | 0.6679 | 0 | 0.1295 |
| HTX_NE_14 | 10 | 0.9547 | 0.8139 | 0.5433 | 0 | 0.6874 |
| HTX_NE_17 | 4 | 0.9752 | 0.781 | 0.4021 | 0 | 0.1114 |
| HTX_NE_8 | 8 | 0.9579 | 0.5651 | 0.8465 | 0.1155 | 0 |
| HTX_NE_30 | 2 | 0.9766 | 0.4653 | 0.4334 | 0 | 0.0354 |
| HTX_NE_32 | 3 | 0.9604 | 0.7891 | 0.8737 | 0.4034 | 0 |
| HTX_NE_18 | 1 | 1 | 0.7433 | 0.9579 | 0.0289 | 0 |
| HTX_NE_21 | 3 | 0.9881 | 0.1866 | 0.7862 | 0.2911 | 0 |
| HTX_NE_22 | 1 | 0.9786 | 0.7421 | 0.6481 | 0.2335 | 0 |
| HTX_NE_27 | 3 | 0.7537 | 0.9108 | 0.6571 | 0 | 0.2047 |
| HTX_NE_2 | 20 | 0.9707 | 0.4281 | 0.5538 | 0 | 0.1207 |
| HTX_NE_4 | 8 | 0.9754 | 0.707 | 0.2352 | 0 | 0.4098 |
| HTX_NE_15 | 8 | 0.958 | 0.777 | 0.7917 | 0 | 0.0479 |
| HTX_NE_16 | 1 | 0.8693 | 0.8303 | 0.5944 | 0 | 0.4253 |
| HTX_NE_6 | 19 | 0.9828 | 0.6671 | 0.9432 | 0.0245 | 0 |
| HTX_NE_29 | 4 | 0.9901 | 0.3656 | 0.6296 | 0 | 0.5179 |
| HTX_NE_31 | 1 | 1 | 0.8964 | 0.8619 | 0 | 0.0276 |
| HTX_NE_19 | 2 | 0.9932 | 0.8115 | 0.8466 | 0.2985 | 0 |
| HTX_NE_20 | 7 | 0.936 | 0.0533 | 0.8795 | 0 | 0.6985 |
| HTX_NE_25 | 4 | 0.9935 | 0.3432 | 0.9882 | 0 | 0.3524 |
| HTX_NE_26 | 6 | 0.9877 | 0.8673 | 0.9136 | 0.0485 | 0 |
| HTX_NE_13 | 15 | 0.6314 | 0.0183 | 0.1661 | 0 | 0.6316 |
| HTX_NE_28 | 20 | 0.9659 | 0.5147 | 0.7258 | 0.0671 | 0 |
| HTX_NE_3 | 19 | 0.9868 | 0.5228 | 0.3832 | 0 | 0.4487 |

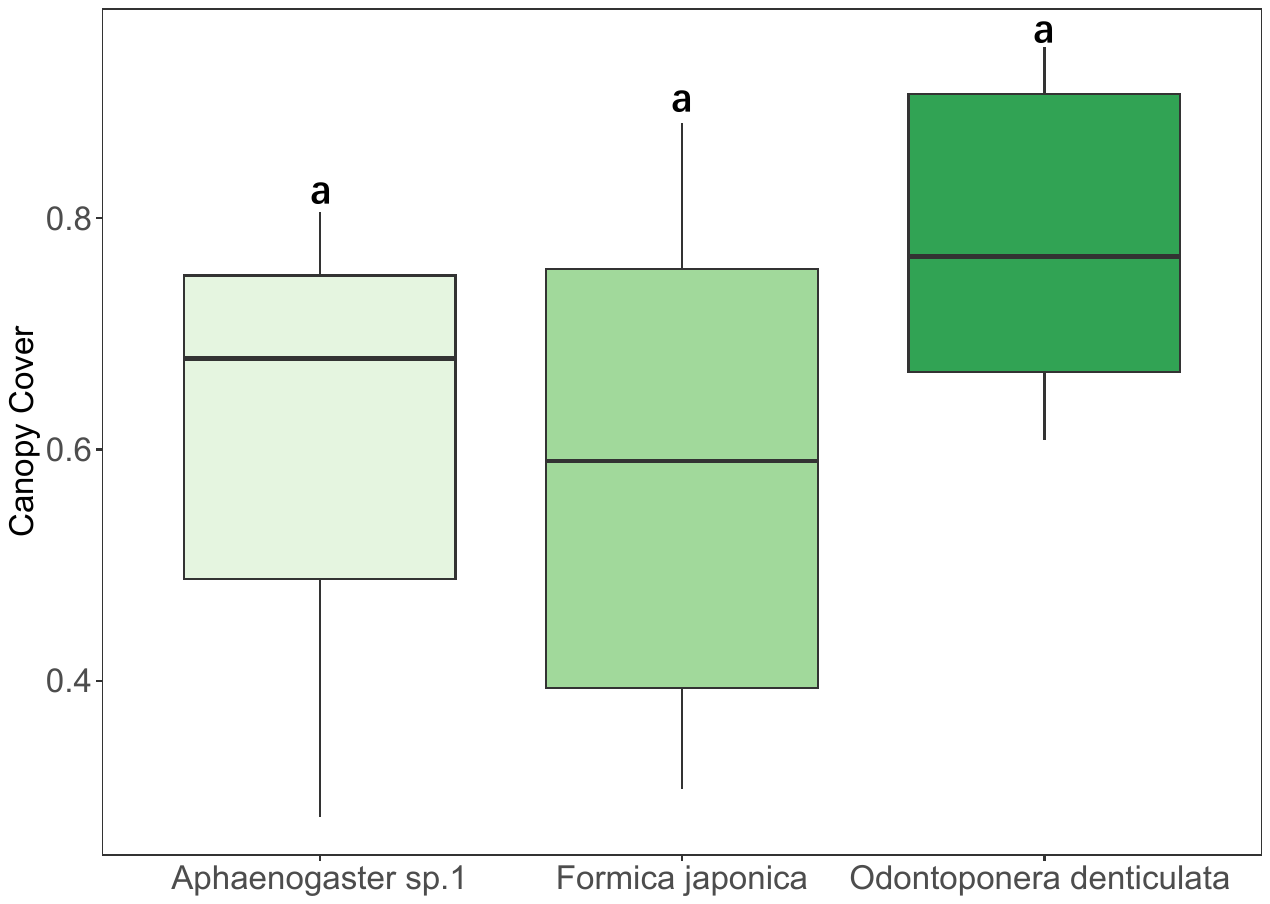

**Figure S1.** Canopy coverage of nest sites of *Aphaenogaster sp.1* (left), *Formica japonica* (middle) and *Odontoponera denticulata* (right).
